# Quantification of GC-biased gene conversion in the human genome

**DOI:** 10.1101/010173

**Authors:** Sylvain Glémin, Peter F. Arndt, Philipp W. Messer, Dmitri Petrov, Nicolas Galtier, Laurent Duret

## Abstract

Many lines of evidence indicate GC-biased gene conversion (gBGC) has a major impact on the evolution of mammalian genomes. However, up to now, this process had not been properly quantified. In principle, the strength of gBGC can be measured from the analysis of derived allele frequency spectra. However, this approach is sensitive to a number of confounding factors. In particular, we show by simulations that the inference is pervasively affected by polymorphism polarization errors, especially at hypermutable sites, and spatial heterogeneity in gBGC strength. Here we propose a new method to quantify gBGC from DAF spectra, incorporating polarization errors and taking spatial heterogeneity into account. This method is very general in that it does not require any prior knowledge about the source of polarization errors and also provides information about mutation patterns. We apply this approach to human polymorphism data from the 1000 genomes project. We show that the strength of gBGC does not differ between hypermutable CpG sites and non-CpG sites, suggesting that in humans gBGC is not caused by the base-excision repair machinery. We further find that the impact of gBGC is concentrated primarily within recombination hotspots: genome-wide, the strength of gBGC is in the nearly neutral area, but 2% of the human genome is subject to strong gBGC, with population-scaled gBGC coefficients above 5. Given that the location of recombination hotspots evolves very rapidly, our analysis predicts that in the long term, a large fraction of the genome is affected by short episodes of strong gBGC.

## Introduction

The process of GC-biased gene conversion (gBGC) has a major impact on the evolution of mammalian genomes (Duret and Galtier 2009; Romiguier et al. 2010; Katzman et al. 2011) and is known or suspected to a play a role in many other groups of eukaryotes (Webster et al. 2006; Escobar et al. 2011; Pessia et al. 2012; Serres-Giardi et al. 2012). gBGC is a recombination-associated process favoring G:C (S for strong, hereafter) over A:T (W for weak, hereafter) bases during the repair of mismatches that occur within heteroduplex DNA during meiotic recombination (Marais 2003; Lesecque et al. 2013). From a population genetics point of view, gBGC is equivalent to natural selection in favor of S alleles, increasing their frequency and probability of fixation (Nagylaki 1983). gBGC therefore tends to increase GC content and W→S substitution rates in highly recombining regions.

There are at least two reasons why we should worry about gBGC. First, as recombination rate is highly heterogeneous across the genome and most recombination events occur in evolutionarily short-lived hotspots (Myers et al. 2005; Ptak et al. 2005; Winckler et al. 2005; Coop and Myers 2007; Auton et al. 2012), gBGC-induced GC-enrichment is expected to occur through short, localized episodic events. Such a sudden, locus and lineage specific acceleration of substitution rates can easily mimic the signature of positive selection (Galtier and Duret 2007; Berglund et al. 2009; Ratnakumar et al. 2010; Kostka et al. 2012). Accordingly, it was estimated that up to 20% of signatures of positive selection in the human genome could be explained by gBGC (Ratnakumar et al. 2010). Clearly, the effects of gBGC must be taken into account seriously in studies of molecular adaptation in humans, mammals and other taxa.

Secondly, gBGC can actually oppose natural selection. This occurs when the S allele is less favorable for the fitness than the W allele. In this case, gBGC tends to maintain deleterious alleles at intermediate or high frequency in populations, possibly until fixation, depending on selection and dominance coefficients (Glémin 2010). Accordingly, gBGC tracts are enriched in disease-associated polymorphisms (Capra et al. 2013) and W→S disease-causing mutations segregate at higher frequency than S→W mutations (Necsulea et al. 2011; Lachance and Tishkoff 2014). High rates of fixation of non-synonymous, likely deleterious, mutations are also associated with gBGC episodes in primates (Galtier et al. 2009).

The magnitude of the above-mentioned effects strongly depends on the intensity of gBGC that can be measured by the population-scaled coefficient *B* = 4*N*_*e*_*b*, where *N*_*e*_ is the effective population size and *b* is the intensity of the conversion bias (Nagylaki 1983). Similar to selection, gBGC is only considered to be effective in that it dominates over random genetic drift if *B* is substantially greater than one. For example, the magnitude of gBGC-induced deleterious effects depends on the distribution of *B* values relative to selection: strong gBGC episodes in a few hotspots is a more harmful situation than homogeneous but low gBGC level (Glémin 2010). For a proper assessment of the impact of gBGC on genome evolution, it is therefore essential to accurately quantify the *B* parameter.

Previous studies have used substitution patterns along phylogenetic lineages to estimate the intensity of gBGC. On average over the whole genome, gBGC was found to be relatively weak *B* = 0.2 to 0.36 (Lachance and Tishkoff 2014). However, based on the estimated proportion of recombination hotspots, Duret and Arndt (2008) evaluated that an average gBGC intensity of *B* = 5 to 6.5 in these hotspots are required to explain the patterns of substitution rates in the human lineage. Recently, Lartillot (2013b) developed a Bayesian method that directly estimates *B* along a phylogeny, incorporating variations both among branches and among genes. Analyzing sets of exons at the scale of the mammalian phylogeny, he showed that *B* could reach average values of about 5 in small-sized mammalian lineages that have high effective population size, with a small percentage of exons evolving under very strong gBGC (*B* > 10). He also confirmed that gBGC is weaker in the human lineage, and more generally in primates than in small-sized, short-lived mammals, which can explain the erosion of GC-rich isochores in this group (Duret et al. 2002; Duret et al. 2006). Capra et al. (2013) also developed a phylogenetic method to capture gBGC heterogeneity and detect gBGC tracts, which they applied to the human and chimp genomes. However, these authors did not quantify the intensity of gBGC in these tracts. In fact, their method requires fixing the value of *B* expected in hotpots (they used *B* = 3). These two methods were successful in capturing (part of) the heterogeneity of gBGC genome-wide, but they describe and quantify the process over millions of years of evolution. Because recombination hotspots, and hence also gBGC hotspots, have a very short lifespan (Ptak et al. 2005; Winckler et al. 2005; Auton et al. 2012; Lesecque et al. 2014) the intensity of gBGC currently experienced by the human population cannot be properly estimated by the methods described above.

Estimates of gBGC in more recent time periods can in principle be obtained from polymorphism data by fitting models of gBGC to the site frequency spectra (SFS) of W→S and S→W mutations (hereafter denoted WS and SW respectively). Within this framework, Spencer et al. (2006) estimated *B* = 1.3 for the 20% highest recombination fraction of the human genome. However, several methodological issues have not been considered in their approach. First, as demography also affects SFS, it must be taken into account into inference approaches. This can be achieved by incorporating a demographic scenario into the model (usually a simple change in population size is used) (Eyre-Walker et al. 2006; Boyko et al. 2008) or by adding noise parameters to account for the non-selective factors that affect the shape of the SFSs (Eyre-Walker et al. 2006 and see below). Second, errors in the polarization of mutations into ancestral and derived alleles, especially because of homoplasy due to CpG hypermutability, are known to affect the SFS, which can lead to spurious signatures of gBGC (Hernandez et al. 2007). One way to circumvent this problem is to use folded spectra, in which mutations are not polarized. However, gBGC intensity can be estimated from the shape of the folded SFS only under the assumption of mutation/gBGC/drift balance equilibrium (Smith and Eyre-Walker 2001). When this assumption is relaxed, derived allele frequency (DAF) spectra are required to disentangle mutation bias and gBGC. Recently, De Maio et al. (2013) combined polymorphism and divergence data in a global framework to both correct for polarization errors due to CpG and distinguish mutation bias from gBGC (De Maio et al. 2013). However, they assumed a constant population size in their model. Finally, a third issue concerns the dynamics of gBGC episodes. Both Spencer et al. (Spencer et al. 2006) and De Maio et al. (2013) found rather low values of gBGC (maximum around *B* = 1) but they did not properly take gBGC heterogeneity into account, and it is not clear how this affects *B* estimates.

Here, we propose a new framework for estimating the intensity of gBGC that solves the issues discussed above. We show by simulations that an important bias in SFS-based estimates of gBGC due to mis-polarization has been consistently overlooked in previous studies, but can be fully corrected within the framework of our method. We also show that strong heterogeneity in *B* can lead to its underestimation, and develop an extension of our approach that accounts for this problem. We apply our inference method to the African (AFR) population of the 1000 genomes dataset (Abecasis et al. 2012) to quantify gBGC and its variation across the human genome and analyze the effect of local recombination rate on these variations.

## Results

### Signatures of gBGC in DAF spectra are obscured by (unexpected) polarization artifacts

To investigate fixation biases affecting WS and SW mutations in the human genome, we analyzed SNP data from the AFR population of the 1000 genomes project (Abecasis et al. 2012). We selected all SNPs located in non-coding regions (i.e., presumably neutrally evolving SNPs) from autosomes. We excluded sex chromosomes to avoid biases due to their specific features – both in terms of mutation pattern and demography. Mutations were polarized using ancestral state predictions based on 4-way multiple alignments (*Homo sapiens, Pan troglodytes, Pongo pygmaeus, Macaca mulatta*) (Paten et al. 2008), which are provided in the original SNP data file (Abecasis et al. 2012). We excluded SNPs for which information about the ancestral state was reported as being unreliable (see methods).

We first focused our analyses on non-CpG SNPs. In agreement with previous reports (Katzman et al. 2011), we observed that, on a genome-wide scale, the DAF spectra of WS SNPs are significantly biased towards higher frequencies compared with the DAF spectra of SW SNPs (Figure 1A). As predicted by the gBGC model, this shift in DAF spectra is much stronger for SNPs located in regions of high recombination (Figure 1C) compared to SNPs located in regions of low recombination (Figure 1B), which is in agreement with previous analyses (Katzman et al. 2011; Lachance and Tishkoff 2014). The difference in mean DAF between WS and SW non-CpG mutations increases steadily with increasing recombination rate, from almost 0 (as expected in absence of gBGC and selection) to about 3.5% (Figure 1D).

**Figure 1:**
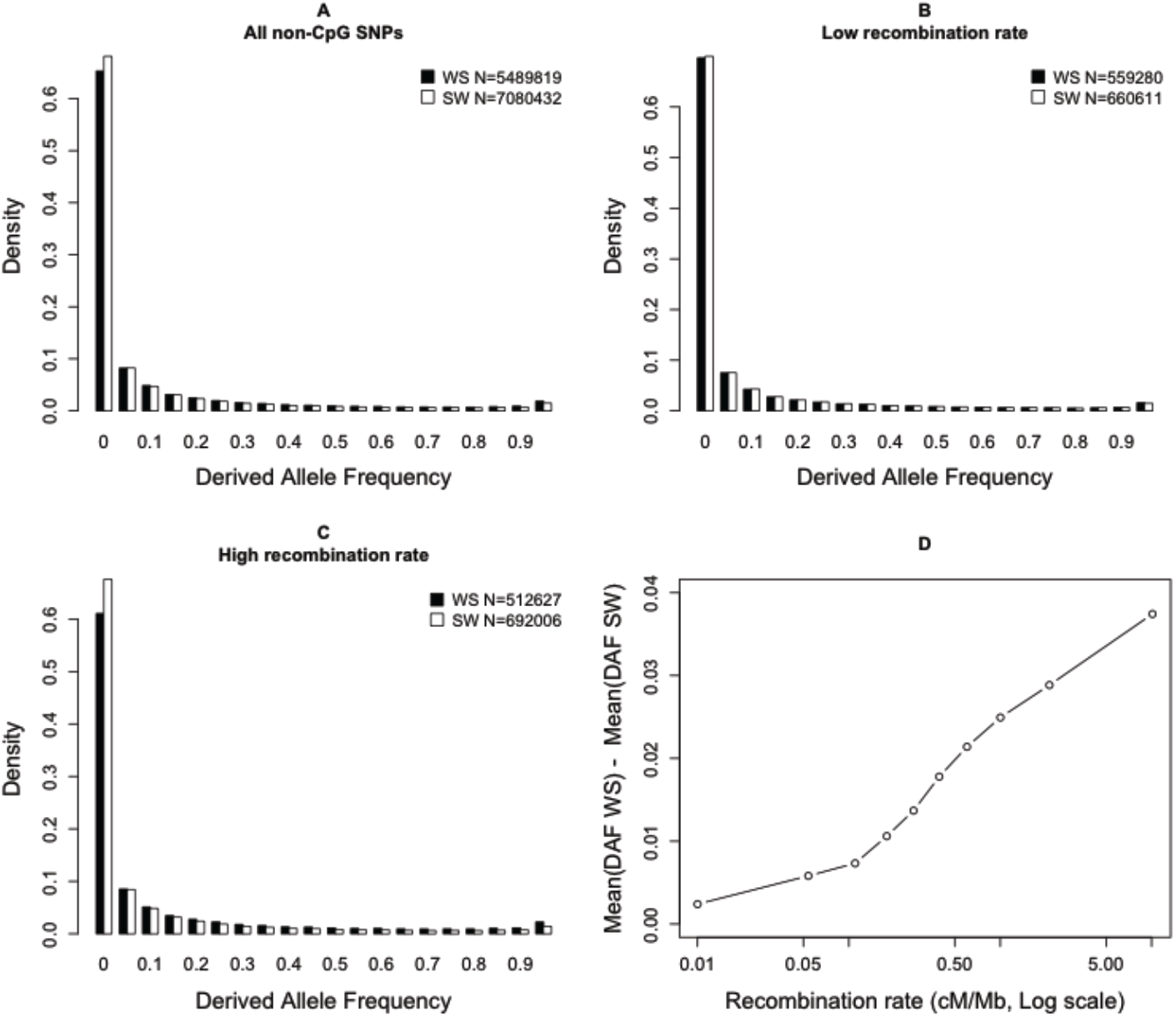
Variations in derived allele frequencies (DAF) according to mutation type (WS or SW) and local recombination rate: non-CpG sites. SNP allele frequencies and polarizations were retrieved from the 1000 genomes phase 1 data set (population panel: AFR). We selected all non-CpG SNPs located in non-coding regions from autosomes. Recombination rates are measured over 5 kb windows centered on each SNP, using HapMap data (Myers et al. 2005). A: DAF spectra of all SNPs. B: DAF spectra of the subset of SNPs located in regions of low recombination (bottom 10%). C: DAF spectra of the subset of SNPs located in regions of high recombination (top 10%). D: differences in mean DAF spectra between WS and SW mutations, according to the local recombination rate. The values in the legends indicate the genome-wide count of SNPs of each category.

The difference in DAF spectra between WS and SW mutations provides information about the intensity of gBGC. We previously developed a generic maximum likelihood model that allows one to quantify the strength of gBGC from the comparison of the DAF spectra of WS and SW mutations, using the DAF spectrum of WW and SS mutations as a neutral reference (2011). In brief, this model can be written as follows. The probability of observing *k*_*i*_ SNPs having *i* derived alleles out of *n* follows a Poisson distribution, *P*(*μ*,*k*_*i*_), with mean:

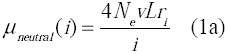

for neutral WW and SS mutations, and

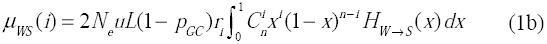

for W→S mutations, and

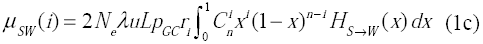

for SW mutations, where *N*_*e*_ is the effective population size, *v* the mutation rate from W to W and from S to S mutations, *u* the mutation rate from W to S, *λu* the mutation rate from S to W, λ being the mutational bias towards AT, *L* the sequence length, and *p*_*GC*_ the GC-content of the sequence. We assumed that *p*_*GC*_ is constant, that is, the ongoing substitution process does not significantly affect base composition over the time scale over which polymorphisms persist. Importantly, this parameterization – instead of assuming a different 4*N*_*e*_*u* for each mutation category – allows estimating the mutational bias, but does not affect the estimate of gBGC parameters. *H*_*SW*_(*x*) and *H*_*WS*_(*x*) are the expected times that a WS, respectively SW, mutation spends at population frequency between *x* and *x* + *dx*. These terms are functions of *B* = 4*N*_*e*_*b*, where *b* is the gBGC coefficient (see Material and Methods and below for the different models we used). The coefficients *r*_*i*_ have been introduced by Eyre-Walker et al. (2006) to account for distortions in DAF spectra due to demography (and/or population structure and/or sampling). The key assumption underlying their approach is that demography affects the DAF spectra of the three different classes of SNPs, neutral, SW and WS, all in a similar way. Thus, the same coefficient *r*_*i*_ is used for each DAF class and corresponds to the deviation from the standard equilibrium model relative to the singleton class, *r*_1_ being set to one.

It has been shown that such models are sufficiently robust to demographic (and/or sampling) (for a detailed discussion of the robustness of this kind of models, see Eyre-Walker et al. 2006; Muyle et al. 2011 and discussion). However, one important difficulty is that the estimation of DAF spectra remains highly sensitive to polarization errors: any WS (respectively SW) mutation observed at frequency *x = i*/*n* in the sample that is mis-polarized is considered as a SW (WS) mutation at frequency (*n* – *i*)/*n*. Given that the majority of derived alleles are rare (i.e., *x* is generally much smaller than 0.5), polarization errors shift the inferred DAF spectra towards higher frequencies. And, given that the SW mutation rate is higher than the rate of WS mutation, the risk of mis-polarization is higher for SW mutations (which are then erroneously counted as WS mutations) (Eyre-Walker 1998). Hence, this polarization artifact leads to overestimating the fixation bias in favor of WS mutations (Hernandez et al. 2007). This artifact is expected to be particularly strong at hypermutable CpG sites, where the inference of the ancestral state is less reliable. And indeed, CpG sites show very peculiar DAF spectra, with a strong peak of WS SNPs segregating at very high frequency (Figure 2A). One possible interpretation is that gBGC might be much stronger on CpG than on non-CpG sites. However this peak is observed regardless of recombination rate (Figure 2B, 2C), and the difference in mean DAF between WS and SW mutations is very high (˜8%) even in regions of very low recombination (Figure 2D). All these observations indicate that the strong excess of WS CpG SNPs segregating at very high frequency is not due to gBGC.

**Figure 2:**
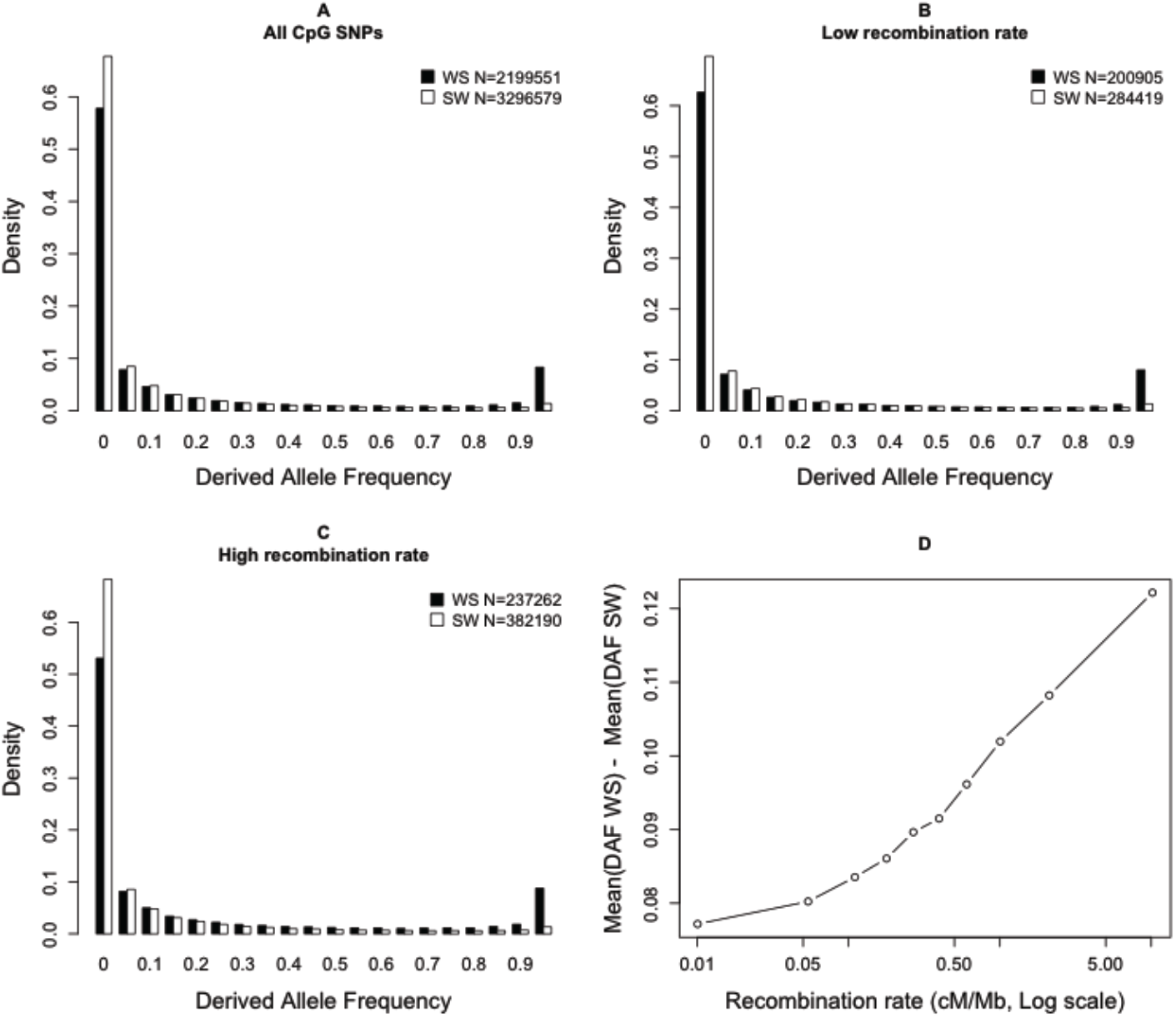
Variations in derived allele frequencies (DAF) according to mutation type (WS or SW) and local recombination rate: CpG sites. Information similar to this given in Figure 1, but here for CpG sites only.

To assess the impact of polarization errors on DAF spectra and on estimators of gBGC strength, we performed extensive simulation analyses (see details in Material and Methods). Simulation parameters were set so as to mimic the situation observed in the human genome, where we estimate that the polarization error rate is about 1% to 4% when using the polarization provided by the 1000 genomes data (see below). In the human genome, as in other mammals, the base composition varies strongly along chromosomes, and generally does not correspond to the mutational equilibrium (Duret and Arndt 2008). We therefore simulated genomes composed of sequences of different GC-content, subject to the same mutational bias (λ= 2). We simulated both genomes with gBGC (with stronger gBGC in regions of higher GC-content) and genomes not subject to any gBGC.

Our simulations revealed both expected and unexpected patterns. As expected, and in agreement with previous reports (Hernandez et al. 2007), gBGC is overestimated when the polarization error rate is higher for SW mutations than for WS mutations (typically as for CpG sites) (Figure S1). However, even when polarization error rates are symmetrical (i.e., we assume the same rate of polarization error for WS and SW mutations), estimates of *B* are biased. This bias leads to a spurious positive relationship between *B* and the local GC-content, and can even lead to the inference of negative average *B* values (Figures 3A and 3B). This surprising result is explained by the fact that, in our simulations, the ratio of WS to SW mutations increases with GC content – this is so because we model the non-equilibrium situation of GC rich regions in humans (see above). The bias is only suppressed when there is an equal number of WS and SW mutations, i.e., when the base composition closely reflects the mutational equilibrium, *p*_GC_ = 1 / (1 + λ). It is therefore crucial to take this bias into account for any method based on DAF spectra that distinguish between WS and SW polymorphisms.

**Figure 3:**
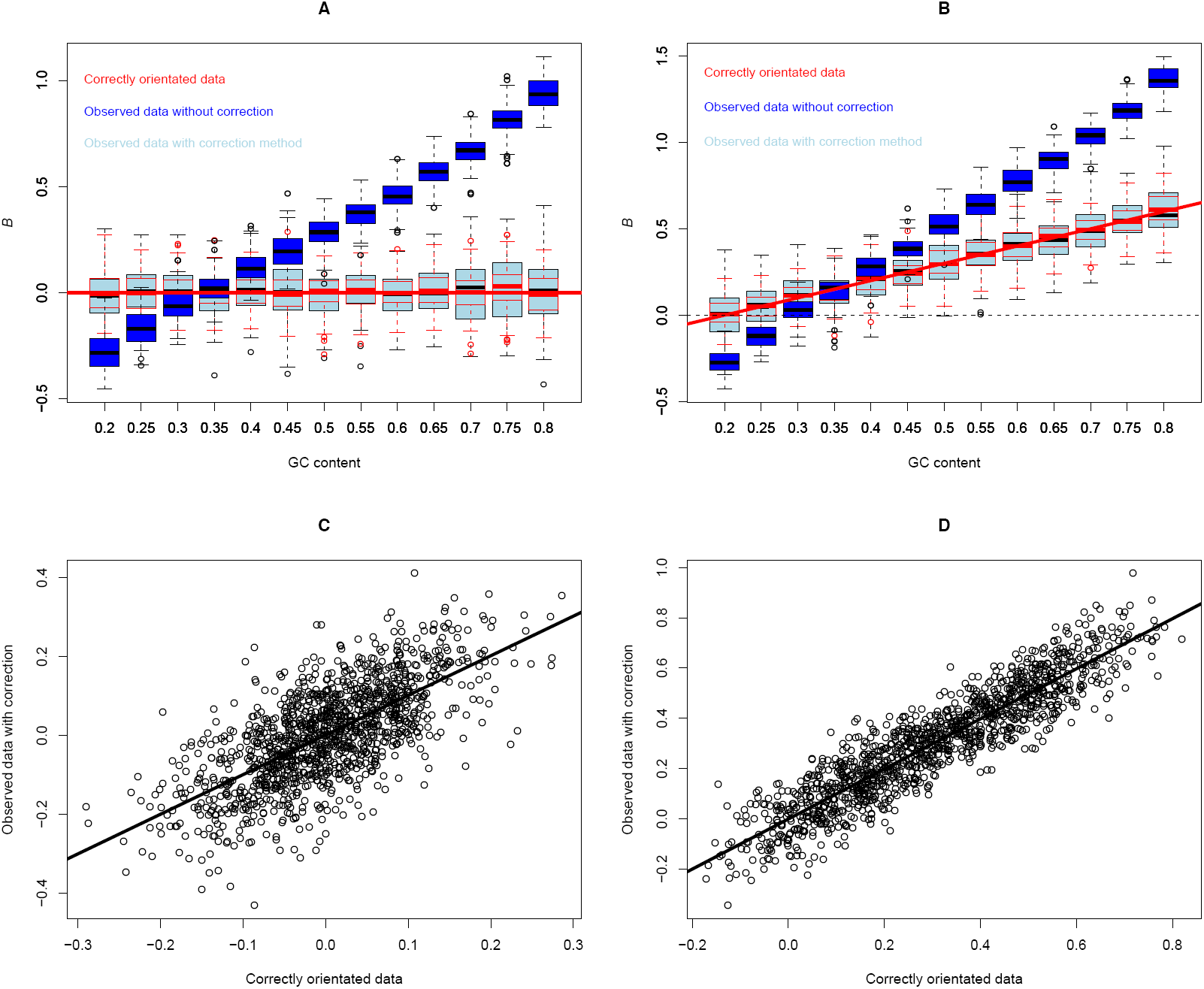
Effect of polarization errors on *B* estimates and accuracy of the correction method. Estimation of *B* as a function of GC content in simulated datasets: *B* = 0 for any GC content (A and C) and *B* linearly increases with GC content (B and D) (red lines in A and B). For a given simulated dataset (= correctly orientated data), polarization errors (*e*_neutral_ = *e*_WS_ = *e*_SW_ = 0.03) were secondarily added (= observed data). *B* values were estimated using the model without error correction (M1, see main text and Table 1) and with error correction (M1*). Box-plots correspond to the *B* estimates of 100 correctly orientated datasets using M1 model (red), 100 observed datasets using M1 model (dark-blue), 100 observed datasets using M1* model (light-blue). Figures C and D show the correlation between *B* values estimated from correctly orientated data with M1 model and *B* values estimated from observed data with the M1* model. The regression line is indistinguishable from the diagonal *y* = *x*.

**Table 1:** Number of parameters for the different models. The saturated model has 3*n* parameters.

| Models | Without error | With error |
| --- | --- | --- |
| M0 | $3 + n - 2$<br>$\theta, \theta_{WS}, \lambda, r_i$ | $6 + n - 2$<br>$\theta, \theta_{WS}, \lambda, e_{neutral}, e_{WS}, e_{SW}, r_i$ |
| M1 | $4 + n - 2$<br>$\theta, \theta_{WS}, \lambda, B, r_i$ | $7 + n - 2$<br>$\theta, \theta_{WS}, \lambda, B, e_{neutral}, e_{WS}, e_{SW}, r_i$ |
| M2a | $5 + n - 2$<br>$\theta, \theta_{WS}, \lambda, B, f, r_i$ | $8 + n - 2$<br>$\theta, \theta_{WS}, \lambda, B, f, e_{neutral}, e_{WS}, e_{SW}, r_i$ |
| M2b | $6 + n - 2$<br>$\theta, \theta_{WS}, \lambda, B_0, B_1, f, r_i$ | $9 + n - 2$<br>$\theta, \theta_{WS}, \lambda, B_0, B_1, f, e_{neutral}, e_{WS}, e_{SW}, r_i$ |
| M3 | $4 + n - 2$<br>$\theta, \theta_{WS}, \lambda, B_0, r_i$ | $7 + n - 2$<br>$\theta, \theta_{WS}, \lambda, B_0, e_{neutral}, e_{WS}, e_{SW}, r_i$ |

### Correcting for polarization error in estimating the intensity of gBGC: a new method

Several methods have been developed to cope with polarization errors, especially to take CpG hyper-mutability into account (Hernandez et al. 2007; Duret and Arndt 2008; De Maio et al. 2013) However, although these methods suppress the bias in the inference of ancestral states, symmetrical polarization errors remain, and our simulations clearly showed that even unbiased mis-polarization is problematic as far as SFS analysis is concerned. Here we propose an alternative approach that incorporates polarization error rates directly into the estimation procedure. The rationale of the method is the same as for the generic model described above, except that, here, the probability of observing *k*_*i*_ SNPs having *i* derived alleles out of *n* follows a Poisson distribution, *P*(μ,*k*_*i*_), with mean:

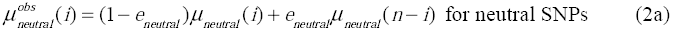

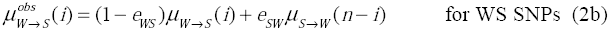

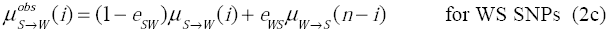

where the “true” *μ* are given by equations (1) and *e*_*neutral*_, *e*_*WS*_ and *e*_*SW*_ are polarization error probabilities, which are estimated jointly with the other parameters of the model. We thus have four possible models: *B* = 0 (M0) and *B* ≠ 0 (M1), without error correction, and the same with error correction (M0* and M1*). The four models can be compared by likelihood ratio test (LRT) with the appropriate degrees freedom (see Table 1). The goodness of fit of these models can then be assessed by comparison with the likelihood of the saturated model, in which every class of each SFS has its own parameter.

We evaluated our new method under different conditions (symmetrical vs. asymmetrical error rates, stable vs. non-equilibrium populations). Simulations show that our method performs well in all tested conditions and accurately recovers the true simulated value of *B* (Figure 3 and Figures S1 and S2). We compared the M1* model applied to datasets with polarization errors to the M1 model applied to the same datasets without errors. The two estimates of *B* were very well correlated with no bias (the regression line was indistinguishable from the *y* = *x* line, Figures 3C and 3D). We also checked the accuracy of the estimation of error rates. These estimates suffer from a large variance. Because error rates are low and bounded to zero, large variance tends to increase error rate estimates on average. As a consequence, the mean estimate of *e*_*WS*_ (resp. *e*_*SW*_) tended to slightly increase (resp. decrease) with GC content. Once again, this is explained by the fact that the number of WS (resp. SW) mutations, and hence the power to estimate *e*_*WS*_ (resp. *e*_*SW*_), decreases (resp. increase) with GC-content (see Figure S3). However, this bias did not affect the estimation of *B*, as shown above.

### A moderate genome-average gBGC intensity in humans

We applied our method on the AFR population of the 1000 genomes dataset (Abecasis et al. 2012), using all non-coding SNPs (whole dataset), only non-CpG SNPs (non-CpG datmutation rate fromaset) and only CpG SNPs (CpG dataset). Several parameters of the model (mutation rates, gBGC strength, polarization error rates) are susceptible to variations along chromosomes. We therefore performed parameter estimations individually on 1 Mb-long windows (non-overlapping) across the genome. Sex chromosomes were excluded from the analysis. For a few windows, one or more models did not converge and these windows were excluded from the analyses. The final numbers of windows for the three datasets are 2669, 2665, and 2644, respectively. Each window was characterized by its average GC-content and recombination rate. Because local GC-content and recombination rates can be different between CpG and non-CpG sites within a given window, for each dataset we computed GC-content at 100bp and recombination rate at 5kb around each SNP and averaged these values over the SNPs of each window.

Over the whole dataset, estimates of *B* obtained with model M1* ranged from -0.70 to 2.06 with a median of 0.35 and a mean of 0.38 (Figures 4). A negative *B* was estimated for only 232 out of 2669 windows and only 11 (5%) were significantly different from 0. In contrast, over the 2437 windows with a positive *B*, 1458 (60%) were significantly different from 0 (Figure 4). As expected, *B* was strongly correlated with the recombination rate (R^2^ = 0.27). We also observed a correlation between *B* and GC content (R^2^ = 0.14). Multiple regression analysis showed that this correlation is essentially due to the known correlation between recombination rate and GC-content (R^2^ = 0.18): the variance of B explained by GC-content and recombination together (R^2^ = 0.30) was only slightly greater than that explained by recombination alone (R^2^ = 0.27). Given that the density in CpG sites increases with GC-content, CpG SNPs are enriched in GC-rich genomic regions and, hence, in regions of high recombination. Consistently, on average *B* was higher for CpG sites compared to non-CpG sites (Table 2). However, for a given recombination rate and local GC content there was virtually no difference in strength of gBGC between CpG and non-CpG sites (Figure 5). Although the effect of the category of sites was still significant after error correction (because of the size of the dataset), it only explained 1% of the variance in *B* when polarization error was correctly accounted for (but 38% otherwise, see Figure 5).

**Figure 4:**
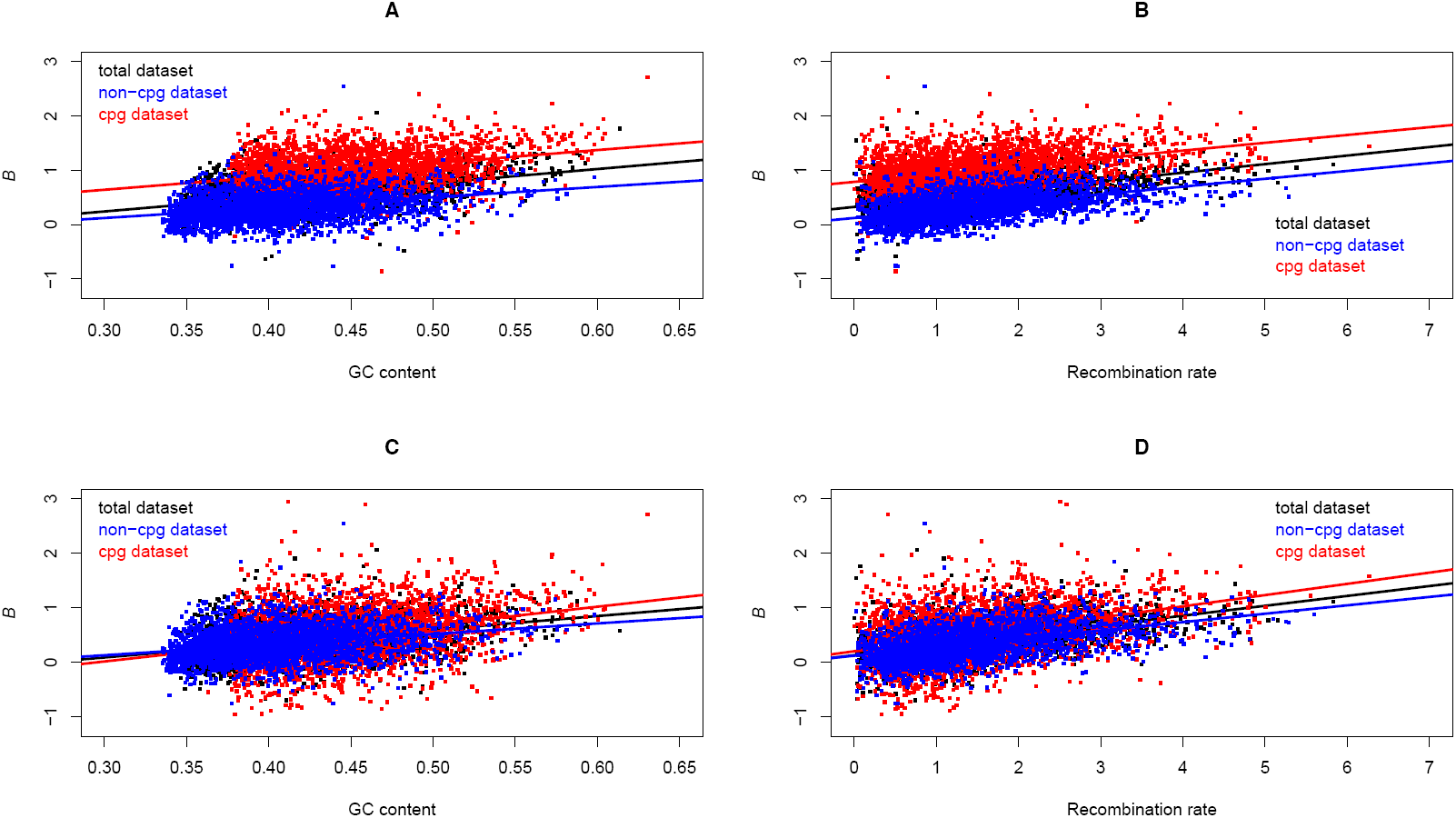
*B* estimates for one-megabase windows as a function of recombination rate and GC content. Values of *B* were estimated with the M1* model. Grey (resp. orange) dots correspond to *B* values non-significantly (resp. significantly) different from 0. The regression lines and the spearman correlation coefficients are given in the plots. ***: p-values < 10^-15^. To be congruent with Figure 5, GC-content was measured over 100bp and recombination rate over 5kb around each SNP and then averaged over each window.

**Table 2:** Estimates of average polarization error rates and gBGC strength (*B*) obtained with model M1* on 1Mb-long genomic windows, and comparison with estimates of *B* obtained without correction for polarization errors (model M1). Polarization error rates: SW mutations (e_*SW*_), WS mutations (e_*WS*_), SS+WW mutations (*e*_*neutral*_).

| Sites | $e_{SW}$ | $e_{WS}$ | $e_{neutral}$ | $B$ | |
| --- | --- | --- | --- | --- | --- |
|  |  |  |  | M1* | M1 |
| All | 1.8% | 1.0% | 0.8% | 0.38 | 0.55 |
| Non-CpG | 0.7% | 1.0% | 0.9% | 0.33 | 0.32 |
| CpG | 4.1% | 0.7% | 0.6% | 0.50 | 1.00 |

**Figure 5:**
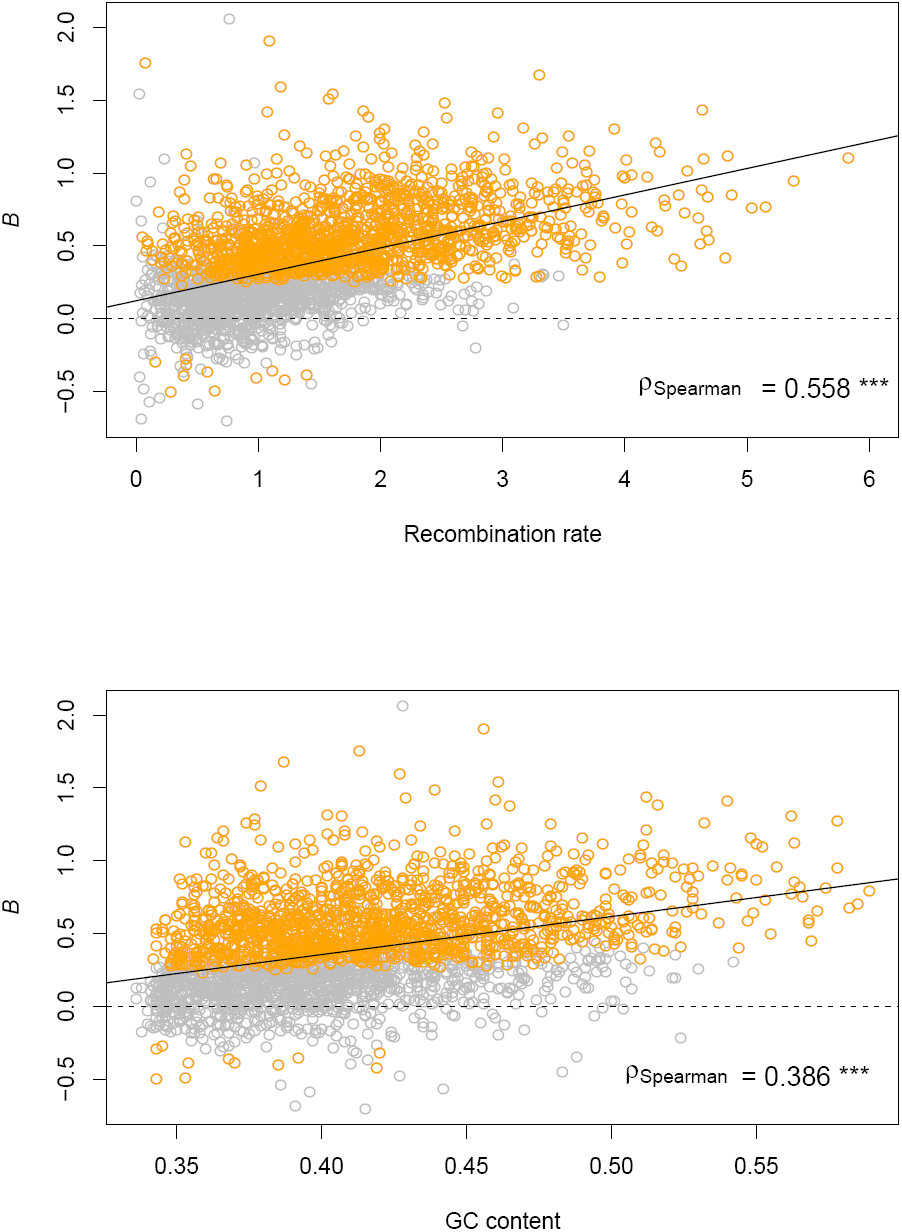
Comparison of the *B* estimated with and without error correction. Values of *B* estimated without error correction (M1 model, A and B) and with error correction (M1* model, C and D) as a function of GC content (A and C) and recombination rate (B and D) for the whole dataset (black), the non-CpG dataset (blue), and the CpG dataset (red). To take into account differences in local GC-content and recombination rates between CpG and non-CpG sites in the same window, we measured GC-content over 100bp and recombination rate over 5kb around each SNP and then averaged them over each window. When the non-CpG and the CpG datasets are analyzed jointly, recombination rate, GC-content, and the category of sites explain, respectively, 11%, 17%, and 38% of the variance in *B* without error correction, and 16%, 4% and 1% with error correction.

Our method estimated an average polarization error rate of about 4% for SW mutations at CpG sites, and 0.6% to 1% for other categories of mutations and sites (Table 2 and Figures S4 to S6). These rates are consistent with the expected rate of homoplasy along the chimpanzee branch, given the branch length between human and chimp. This shows that the method accurately estimates error probabilities, on average, despite the fact that no prior information was included in the model. As predicted by simulations, there was a slight effect of GC content on error estimates: the variance and the mean of *e*_*WS*_ increased with GC content, the variance of *e*_*SW*_ decreased with GC content, while there was no effect of GC content on *e* (Figure S4). Although these error rates are relatively low, they have a strong impact on the quantification of gBGC: on the whole dataset, when ignoring polarization errors *B* is overestimated by 49%, and this overestimate reached 96% for CpG sites (Table 2). Importantly, the difference in estimates of *B* between CpG and non-CpG sites disappeared when polarization errors were accounted for (Figure 4). As predicted by simulations, the correlation between *B* and GC content was lower when error correction was applied (R^2^ = 0.14) than without correction (R^2^ = 0.21). On the contrary, error correction did not affect the correlation between *B* and recombination rate (R^2^ = 0.27 with correction vs. R^2^ = 0.29 without correction). Our method thus appears efficient to correct for biases induced by GC-content dependent polarization errors at both CpG and non-CPG sites. In what follows, all results are presented for the whole dataset (CpG + non-CpG) with correction for mis-orientation of SNPs.

### gBGC is underestimated when its strength varies along a chromosome

In agreement with previous studies (Spencer et al. 2006; De Maio et al. 2013), our genome-wide estimates of *B* are relatively low, in the nearly-neutral area. At first sight, this appears to be in contradiction with other analyses reporting episodes of very strong gBGC (Galtier and Duret 2007; Ratnakumar et al. 2010). However, the model we used above assumes that all sites in a given window evolve under the same gBGC regime. We thus performed additional simulations to test the robustness of our approach to spatially heterogeneous levels of gBGC. We modeled recombination/gBGC hotspots by considering two categories of SNPs: a fraction, *f*, of SNPs was affected by recombination hotspots with mean gBGC *B*_1_, whereas the other fraction, 1 – *f*, was affected by a basal gBGC level *B*_0_, with 0 ≤ *B*_0_ < *B*_1_. We fixed *B*_0_ and we let *B*_1_ vary to simulate variation in hotspot intensities. For simplicity reason, here we did not include polarization errors in the simulation, neither in estimations. Under this model, the average *B* is equal to (1 – *f*)*B*_0_ + *f B*_1_ and increases linearly with *B*_1_. Contrary to this expectation, we observed that the estimated *B* quickly saturated as *B*_1_ increased (Figure 6A). gBGC is thus underestimated by model M1 when its strength is highly heterogeneous along the chromosome.

**Figure 6:**
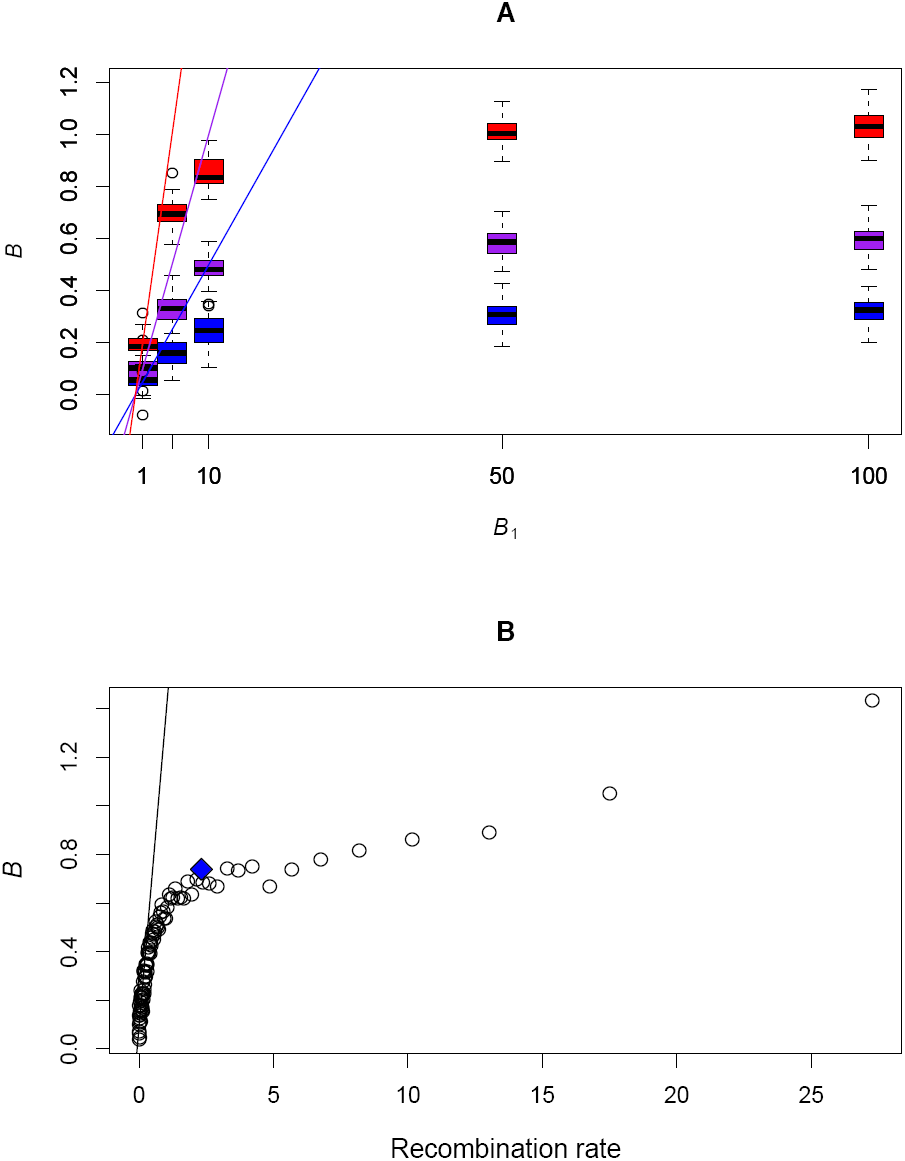
Effect of hotspots on *B* estimates. A) Simulations were performed under model M2b with *B*_0_= 0, *f* = 0.05 (blue), 0.1 (purple) and 0.2 (red). For each *B*_1_ value (x-axis) 100 simulations were performed and the M1 model was applied to estimate B. The lines correspond to the expectation *B* = (1 – *f*)*B*_0_ +*fB*_1_. Very similar results were obtained for *B*_0_= 0.25, and *B*_0_= 0.5 (not shown). B) The whole dataset was divided into centiles of recombination rates computed over 5kb around each SNP. The line corresponds to the regression performed on centiles for which recombination rate was lower than 0.1 cM/Mb. Dots correspond to B estimates under the M1* model. The blue diamond corresponds to *B* estimated using the SNPs belonging to the gBGC tracts detected by Capra et al. (2013).

To check this prediction, we analyzed the human AFR data set in a distinct way: rather than using genomic windows, we grouped SNPs into centiles of local recombination rate (measured on 5kb windows centered on SNPs), thus maximizing the range of expected gBGC intensities among groups of SNPs. As predicted by simulations, the estimated *B* did not increase linearly but roughly log-linearly with recombination rate (Figure 6B). We thus did not estimate very high *B* values, even for the highest recombination rate centiles: the maximum was only *B* = 1.47. This suggests that gBGC is too heterogeneous to be accurately estimated by the simple constant gBGC model (M1a*), even when SNPs are grouped by similar recombination rates.

In order to try to capture this heterogeneity, we introduced two additional models. We first considered a model where gBGC (*B*_1_) only affects a fraction, *f*, of sites, the remaining fraction of sites evolving neutrally (*B*_0_ = 0) (M2a and M2a* with error correction, hereafter). Then we considered a low but non-null basal intensity of gBGC (*B*_0_) for a fraction, 1 – *f*, of sites, and higher gBGC intensity (*B*_1_ > *B*_0_) for the remaining fraction, *f* (M2b and M2b* with error correction, hereafter). Simulations showed that small values of *f* are estimated with large variance, especially when *B*_1_ is small (Figure S7). Moreover, large *B*_1_ are also estimated with large variance (Figure S8). This is due to the fact that above a given threshold (*B* > 20), all values of *B* are expected to give very similar DAF spectra (Figure S9). It is therefore difficult to jointly estimate *f* and *B*_1_ accurately. In the M2b model, *B*_0_ is well estimated, except when *B*_1_ is small (Figure S9). Overall, the M2a model appears more robust than M2b. However, if *B*_0_ > 0, *f* is overestimated and *B*_1_ is underestimated under M2a (see Figures S7 and S8).

To circumvent this difficulty, we used external information to constrain the model (noted M3*). M3* is a version of M2b* in which *f* is fixed to the fraction of recombination hotspots detected by HapMap in each window, and *B*_1_ is set to *ρB_0_*, where *ρ* is the ratio of recombination rates measured in and outside hotspots (see Material and Method). M3* therefore includes a single free gBGC parameter but still allows taking gBGC heterogeneity into account. Applying this model to the human AFR data set, we estimated the distribution of *B* outside (*B*_0_) and within hotspots (*B*_1_) across 2625 one-megabase windows (Figure 7). Excluding the 1% most extreme values, gBGC intensities ranged from -0.22 to 0.98 with a median of 0.16 and a mean of 0.21 outside hotspots, and from -5.66 to 34.23 with a median of 3.55 and a mean of 4.86 within hotspots. Averaging over hotspots and coldspots, the mean *B* equaled 0.52, which is 37% higher than the mean estimated with model M1* (mean *B* = 0.38, Table1). The more negative extreme values within than outside hotspots are simply explained by the constraint that *B*_1_ = *ρB*_0_. Overall, 891 of 2625 windows exhibited values of *B*_1_ higher than 5. Given that hotspots cover on average 6.7% of each window, this indicates that about 2.3% of the genome experience a gBGC intensity higher than *B* = 5.

**Figure 7:**
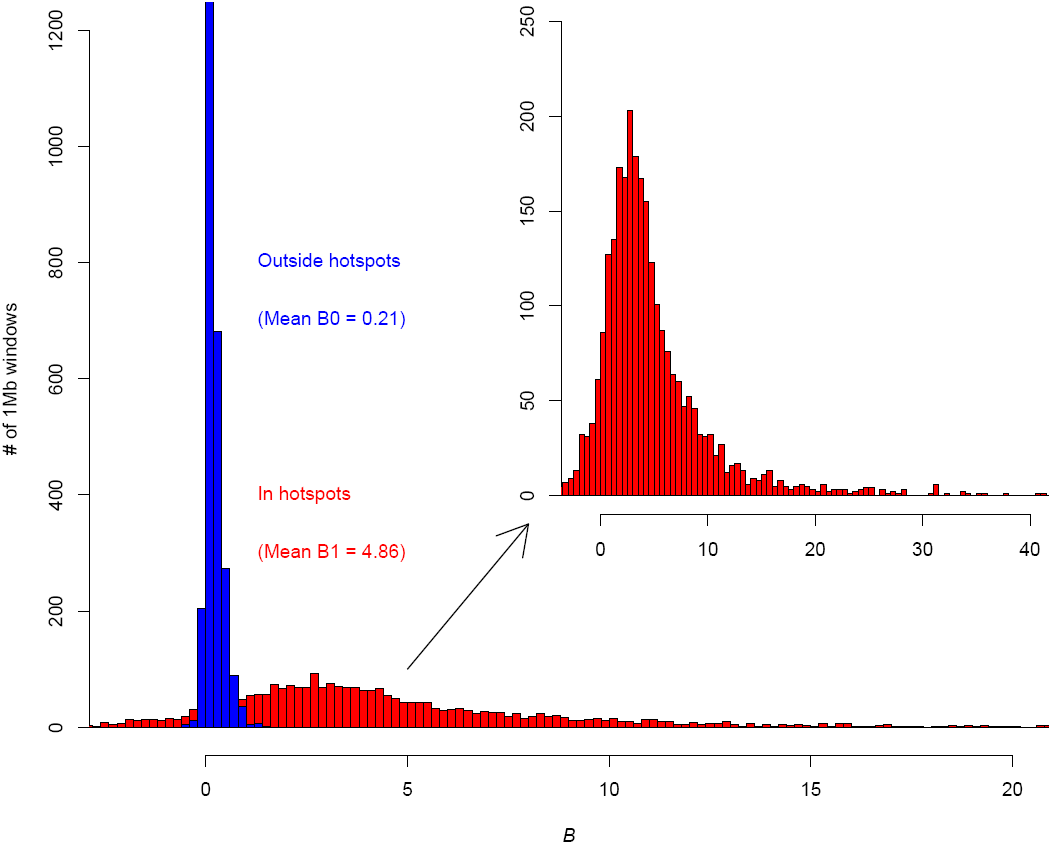
Distribution of *B* in and outside of recombination hotspots for one-megabase windows. Distribution of the estimate of *B*_0_ (= Outside hotpsots, blue) and *B*_1_ (= In hotspots, red) for each one-megabase window. The M2b* model was used and *f*, the fraction of hotspots, was fixed to the observed fraction. The ratio *ρ* = *B*_1_/*B*_0_ was also constrained to be proportional to the ratio of recombination rates in and outside hotspots. Over the 2546 windows, 28 windows were excluded for which at least one of the two estimates of recombination rates was negative. In the inset, the distribution of *B*_1_ is magnified and extended to the whole range of values.

### Mutational bias and global mutational disequilibrium in the human genome

Finally, our method also allows estimating the mutational bias towards AT bases and provides insights into the mutational process genome-wide. Using the M1* model on the whole dataset, we obtained the mean mutational bias (λ) across the genome to be *λ* = 2.08, which is very close to the direct estimate obtained by Kong et al. (Kong et al. 2012) (Figure 8A). Assuming mutational equilibrium, the expected GC-content distribution predicted by the distribution of values of *λ* over analysis windows should be much narrower than the extant distribution with a much lower median (0.33 vs 0.40 see Figure 8B). This striking observation highlights the genome-wide effect of gBGC. However, the two distributions partly overlap, suggesting that some regions of the genome (at the Mb scale) could be at mutational equilibrium. To test for this prediction we show in Text S1 that some insights can be gained by a simple property of folded SFSs, in which SNPs are not polarized. In populations at equilibrium (all *r*_*i*_ = 1) the folded spectrum is symmetrical if and only if GC content is equal to the mutational equilibrium: *p*_GC_ = 1 / (1 + λ). This is true whatever the distribution of *B* and is quite robust to the departure from the demographic equilibrium (*r*_*i*_ ≠ 1). The skewedness of the folded SFS, noted *λ*, is thus a measure of the departure of GC content from its mutational equilibrium: *γ* < 0 (resp. *γ* > 0) indicates that GC content is higher (resp. lower) than its equilibrium. We computed the skewedness of the folded SFS for each window of the whole dataset. Most of the genome has negative skewedness, indicating higher GC content than expected under mutational equilibrium (Figure 8C). Interestingly, skewedness decreases linearly with GC content and extrapolation of the regression line for zero skewedness leads to a mutational equilibrium GC-content of 0.32, which is very close to the value directly estimated from mutation bias (see Figures 8A and 8C).

**Figure 8:**
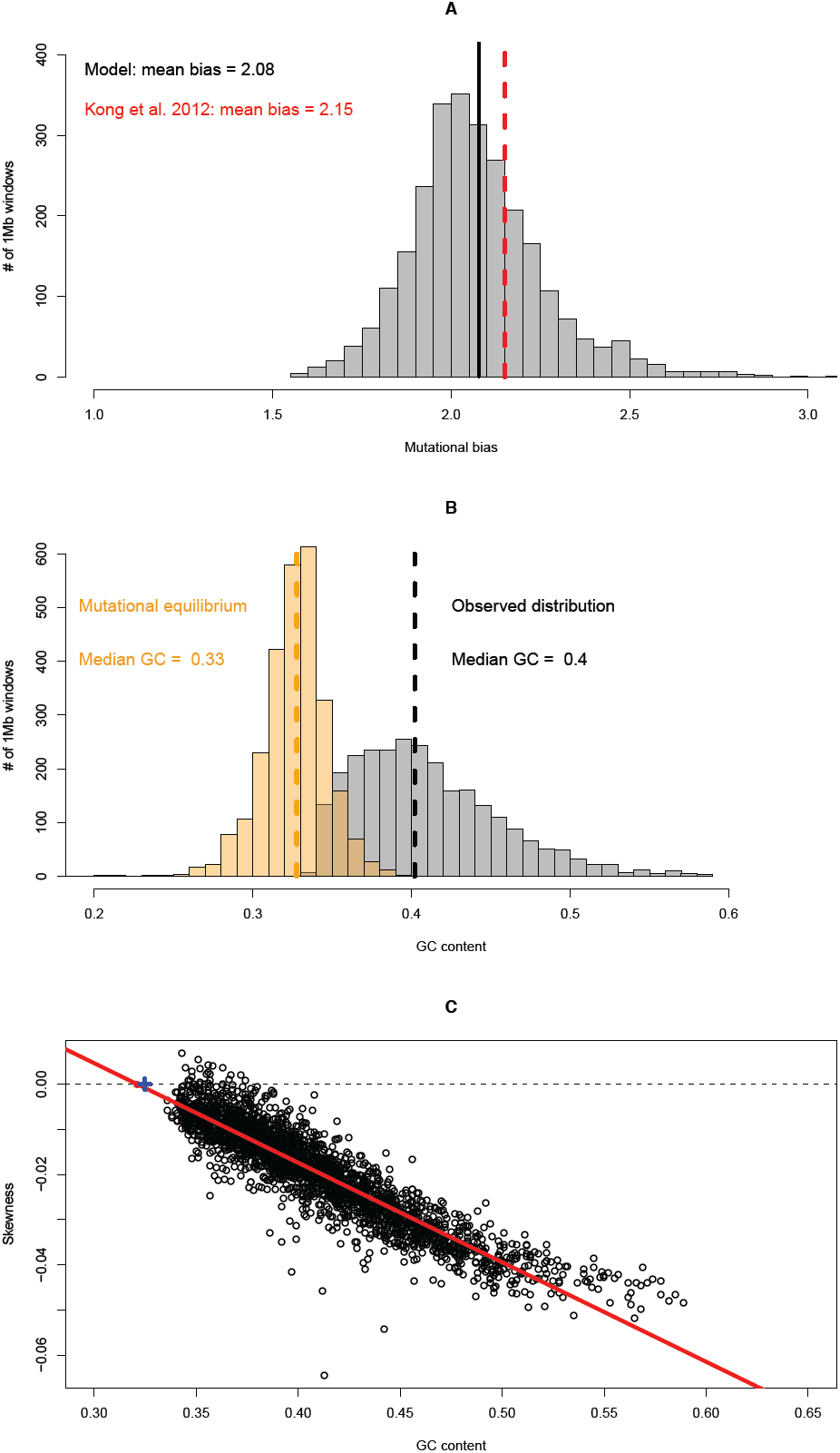
Mutational bias towards AT. A) Distribution of the mutational bias for one-megabase windows.
The distribution of λ was obtained from the M1* model over the 2675 windows. B) Comparison of the observed GC-content distribution and the expected mutational equilibrium distribution. The observed distribution was computed over the 2671 windows, and the expected distribution using the distribution of λ(A) and the relationship: *p*_GC_ = 1 / (1+ λ). C) Skewedness of the folded frequency spectrum for one-megabase windows as a function of GC content. The red line is the regression line. The point for which the skewedness is zero indicates the expected mutational equilibrium GC content: 0.32. The blue cross corresponds to the mutational equilibrium estimated using the mean value of λ(GC = 0.33).

## Discussion

Many lines of evidence show that gBGC is a major determinant of the evolution of GC content in mammalian genomes. Quantifying its intensity throughout the genome is necessary to appreciate its evolutionary and functional impact. As gBGC is driven largely by recombination, which is highly heterogeneous along the genome and episodic in time (Myers et al. 2005; Ptak et al. 2005; Winckler et al. 2005; Coop and Myers 2007; Auton et al. 2012), it is especially important to obtain estimates over short genomic scales and short time scales. So far, such quantifications were still lacking. To achieve this goal we used sequence polymorphism data and tackled several issues associated with the use of such kinds of data. We proposed a new efficient method and provided a fine description of the heterogeneity of the gBGC process along the human genome.

### Methodological issues

DAF spectra potentially contain information about the gBGC process and, more generally, about selection-like processes. However, to correctly infer the intensity of gBGC, two issues need to be addressed: the effect of demography and/or sampling on spectra and the problem of polarization errors. Two alternatives have been proposed to correct for demographic effects. Demographic parameters can be imposed to the estimation model (Boyko et al. 2008) or jointly inferred with selection/gBGC parameters (Keightley and Eyre-Walker 2007). Eyre-Walker et al. (2006) proposed to correct for demography by adding correction parameters for each frequency category. This latter approach is more general because it is valid for any scenario, including specific sampling schemes, which cannot be easily modeled by a simple change in population size. However, it assumes that distortions from the equilibrium expectation are the same for neutral and selected spectra, which should be accurate for weak selection but not for strong selection. Because gBGC is relatively weak globally, it is fully justified to use the second approach, which makes our method quite general and practical for many conditions.

The most serious issue is the spurious signature of gBGC created by polarization errors (Hernandez et al. 2007). Contrary to previous approaches that seek to get accurate reconstruction of ancestral states before applying an inference model, we proposed to include polarization errors directly in the inference model and to estimate them jointly with the other parameters of interest. The advantage of this approach is that it is blind to the underlying process creating polarization errors. It therefore does not require a priori information about processes of sequence evolution, such as a context-dependent mutation rates that take CpG hypermutability into account (Hernandez et al. 2007; Duret and Arndt 2008) Moreover, we showed by simulations that simply correcting the polarization bias between WS and SW mutations is not sufficient because even symmetrical error rates can be problematic (Figure 3).

Overall, we showed by simulations that our joint-inference method performed well under various scenarios. Practically, we also showed that the method corrected well for CpG effects: we observed a clear difference between CpG and non-CpG sites with the basic model without polarization errors, whereas this difference disappeared when we used the model with error correction (Figure 4).

For non-CpG sites, the correction for polarization errors did not affect the estimate of *B* (Table 2). One might therefore argue that the simplest option to avoid biases due to polarization errors consists in excluding CpG sites from the analysis. However, an important drawback of this option is that CpG sites are not uniformly distributed along the genome: the exclusion of CpG sites therefore leads to biases in the sampling towards SNPs located in GC-poor regions, where the recombination rate is on average lower, and thus gBGC is weaker. Hence, to obtain an unbiased estimate of gBGC strength across the entire genome, it is necessary to analyze all categories of SNPs. Moreover, the quantification of gBGC at CpG sites is also interesting in itself for understanding the molecular mechanisms causing gBGC (see below).

Finally, we showed that the strong heterogeneity of the gBGC process made its accurate quantification difficult. On average, the signature of gBGC is weakened by heterogeneity. We thus extended the constant gBGC model to take recombination/gBGC hotspots into account, taking advantage of the detailed knowledge of the recombination landscape in humans that we used to constrain the model and limit the variance on estimates.

It is important to note that the location of recombination hotspots evolves very rapidly. Notably we have shown that human recombination hotspots are at most 0.7 to 1.3 Myrs old (Lesecque et al. 2014). It is therefore likely that DAF spectra at sites that correspond to previous recombination hotspots that are no longer active still retain the hallmarks of past gBGC activity. And conversely, DAF spectra at human recombination hotspots are probably not yet at mutation/drift/gBGC equilibrium. This is why the strength of gBGC cannot be estimated simply by analyzing DAF spectra at presently-active recombination hotspots. Here we modeled hotspot dynamics by considering DAF spectra as a mixture of two categories of sites, supposed to evolve under a stationary regime, which is mathematically convenient. This is clearly an over-simplification, and we suspect that signature of gBGC is also weakened because gBGC is episodic. In the future, a challenging perspective to better quantify the heterogeneity of the gBGC process would be to develop non-stationary models taking into account both heterogeneity between sites and short-lived episodes.

Despite the limitations mentioned above, we suggest that our method can be applied to a broad set of organisms and datasets because a specific knowledge of the demographic history is not required and the effect of polarization errors can be easily corrected for.

### No difference in gBGC strength between CpG and non-CpG sites

The fact that we observed no difference in the strength of gBGC between CpG and non-CpG sites (Figure 4) provides insights about the molecular mechanisms causing gBGC in humans. It is known that the methylation of cytosines at CpG sites is responsible for their hypermutability: the spontaneous deamination of 5-methylcytosine causes the formation of G/T mismatches in DNA that, if not repaired, lead to G:C **→** A:T mutations in the next round of DNA replication. The base excision repair system (BER) plays a major role in the repair of such mismatches. This pathway is initiated by the activity of DNA glycosylases that recognize the G/T mismatch and specifically excise thymines. The resulting gap is ultimately repaired into a G:C base pair (for review, see Sjolund et al. 2013). Mammalian cells possess four enzymes with thymine glycosylase activity (Sjolund et al. 2013). Two of these thymine glycosylases act preferentially at CpG dinucleotides, presumably to limit the hypermutability of these sites: Methyl-CpG Domain Protein 4 (MBD4) and Thymine DNA Glycosylase (TDG) (Sjolund et al. 2013).

Given that the repair of G/T mismatches by BER is systematically directed towards G:C base pairs, it has been hypothesized that this process might be responsible for gBGC in mammals (Brown and Jiricny 1987; Birdsell 2002; Duret et al. 2002; Marais 2003). If this were indeed the case, given the preferential activity of BER at CpG sites, one would then expect a stronger gBGC on CpG than on non-CpG sites. The fact that we do not observe such a pattern strongly argues against this hypothesis. This observation is in accordance with recent results demonstrating that in yeast, gBGC is not caused by BER (Lesecque et al. 2013). The prominent repair pathway during recombination is the mismatch repair (MMR) system (Surtees et al. 2004). In yeast, the analysis of gene conversion tracts indicates that gBGC is most probably caused by MMR (Lesecque et al. 2013). Our observations suggest that this might also be the case in humans.

### Intensity and dynamics of gBGC across the human genome

In agreement with previous studies (Spencer et al. 2006; Capra et al. 2013; Lartillot 2013b; Lachance and Tishkoff 2014) we found that gBGC is weak on average (*B* = 0.52 by averaging M3* estimates over hotspots and coldspots), but widespread along the human genome, which is sufficient to explain that GC content is higher than the expected mutational equilibrium in most regions of the genome (Figure 8). However, average values mask the strong heterogeneity we detected. In highly recombining hotspots, gBGC values can reach high values (*B* > 10, Figure 7) and we evaluated that more than 2% of the genome experience gBGC higher than *B* = 5. Given that the location of hotspots evolves continually (Myers et al. 2005; Ptak et al. 2005; Winckler et al. 2005; Coop and Myers 2007; Auton et al. 2012), this implies that over the long term this process affects a large fraction of the genome.

Previous attempts to quantify the impact of gBGC were based on the analysis of substitution patterns along the phylogeny (Capra et al. 2013; Lartillot 2013b). Capra and colleagues (2013) estimated that about 0.3% of the human genome have been subject to strong gBGC episodes since the divergence from chimpanzee, whereas Lartillot (2013b) did not detect any signature of strong gBGC episodes in primates. This contrast with our results, which indicate that 2% of our genome is currently subject to strong gBGC (*B*>5). The discrepancy is probably due to the fact that these phylogenetic approaches tend to effectively average processes over periods of time (divergence between species) that are much longer than the lifespan of recombination hotspots. Hence, only extremely strong or long-lasting gBGC episodes can be detected by such methods. For a comparison, the distribution of *B* values obtained under the M3* model (Figure 7) indicate that the 0.3% of the human genome with the strongest gBGC, experience *B* values higher than 13.9 (with a mean of 21.6).

Our results also allow us to elucidate the dynamics of gBGC hotspots. If the gBGC tracts detected along the genome by Capra et al. (2013) were still active gBGC hotspots, we should observe high *B* values in these tracts. To test this, we retrieved all SNPs belonging to these tracts on (http://genome-mirror.bscb.cornell.edu) and applied the M1* model. The value we obtained, *B* = 0.74, is higher than the mean computed over the one-megabase windows (*B* = 0.38 with M1* and 0.52 with M3*), but still rather low. Accordingly, the current average recombination rate around these tracts (2.32 cM/Mb) is higher than the genomic mean (1.42 cM/Mb), but does not reach the most extreme values (Figure 6B). These observations suggest that, on average, gBGC is currently not extremely active in these tracts. Thus, most of these tracts probably correspond to ancient recombination hotspots that are no longer active. This is in agreement with the recent findings that current human hotspots are less than 0.7 to 1.3 Myrs old (Lesecque et al. 2014), i.e. much younger than the human-chimpanzee divergence time (7 to 13 Myrs, Langergraber et al. 2012).

### Consequences of transient strong gBGC episodes

As already suspected, our results show that strong gBGC episodes transiently occur along the genome. The consequences of a highly heterogeneous vs. a homogeneous gBGC process are strikingly different even when the mean effect in both scenarios is the same. First, strong gBGC episodes are required to explain substitution hotspots (Dreszer et al. 2007; Kostka et al. 2012; Clement and Arndt 2013) and spurious signature of positive selection (Galtier and Duret 2007; Ratnakumar et al. 2010), but previous studies so far only provided rather low average estimates with maximum *B* values slightly higher than one (Spencer et al. 2006; De Maio et al. 2013; Lartillot 2013b). Here, we directly show that the intensity of gBGC can locally reach values higher than *B* = 5 and even of the order of *B* = 20, which is largely sufficient to explain substitution hotspots. Beyond these technical consequences for the interpretation of genomic patterns, gBGC can counteract selection (Galtier et al. 2009; Lartillot 2013a) and have deleterious consequences (Necsulea et al. 2011; Capra et al. 2013; Lachance and Tishkoff 2014), and it has been shown that strong gBGC episodes in few hotspots have worse consequences than low gBGC levels that are homogeneous along the a chromosome (Glémin 2010). gBGC can thus contribute significantly to the genetic load experienced by human populations. Even though the load a population can tolerate can be high under soft and/or stabilizing selection (Lesecque et al. 2012; Charlesworth 2013), our estimates are quantitatively compatible with potential pathological implications of gBGC as previously proposed (Galtier et al. 2009; Necsulea et al. 2011; Capra et al. 2013; Lachance and Tishkoff 2014).

## Material and methods

### Dataset

We downloaded the 1000 genomes project polymorphism data set (phase 1) (Frazer et al. 2007) from the EBI web site: ftp://ftp.1000genomes.ebi.ac.uk/vol1/ftp/technical/working/20120316_phase1_integrated_release_version2/

This file contains 38,248,808 SNPs, called from the genomes sequences of 1092 individuals from 14 populations including Europe (EUR), East Asia (EAS), sub-Saharan Africa (AFR) and the Americas (AMR). This file also includes predictions of ancestral allele states, inferred from 4-way EPO multiple alignments (*Homo sapiens, Pan troglodytes, Pongo pygmaeus, Macaca mulatta*) (Paten et al. 2008). Details about the procedure used by the 1000 genomes project to infer ancestral states is available here: ftp://ftp.1000genomes.ebi.ac.uk/vol1/ftp/pilot_data/technical/reference/ancestral_alignments/

### README

In brief, the authors first reconstructed ancestral sequences at the two internal nodes of the 4-species phylogeny (using the probabilistic method *Ortheus* (Paten et al. 2008)). Then they retained human-chimpanzee ancestral state predictions that involved no more than one change along the chimpanzee and orangutan lineages. Ancestral state predictions are available for 36,701,805 SNPs (96%). We further excluded 2,606,317 SNPs for which the inference of the ancestral allele was reported as being less reliable (indicated by a lower case in the original file). It should be noted that the reconstruction of ancestral states by the *Ortheus* method does not take into account the hypermutability of CpG sites.

To measure recombination rates we used genetic maps from the HapMap Phase 2 project (Frazer et al. 2007). As these maps are reported in the version hg18 of the human genome assembly, we converted the location of SNPs from hg19 to hg18 coordinates using the liftOver tool (https://genome.ucsc.edu/cgi-bin/hgLiftOver). A small fraction of SNPs (N=31,056) could not be mapped onto hg18 and the SNPs were discarded.

## Maximum-likelihood framework to estimate the intensity of gBGC from site frequency spectra (SFS)

We fitted population genetic models to the derived allele frequency (DAF) spectra to estimate *B* using a maximum likelihood framework similar to Muyle et al. (2011). The generic model is given by equations (1) in the main text. In equations (1), the first term within the integral corresponds to the binomial sampling of *i* alleles in a sample of size *n* given true population-frequency *x*. When *n* is high we can use the continuous approximation that gives very similar results and speeds up and numerical computations:

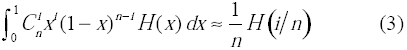

For each subpopulation of the 1000 genomes dataset the frequencies are given in 1/100 so that we set *n* = 100.

We used the following nested models:

**M0: no gBGC:**

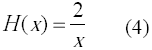

**M1: constant gBGC of intensity *B*** = **4*N*_*e*_*b***:

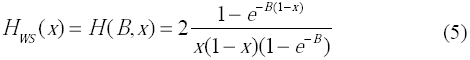

and *H*_SW_(*X*) = *H*(-*B,x*). *B* can be either positive or negative.

**M2a: gBGC hotspots of intensity *B*** = **4*N*_*e*_*b* in frequency *f***:

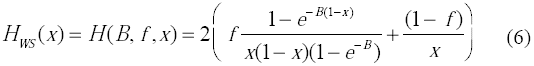

and *H*_SW_(*X*) = *H*(-*B, f,x*). *B* can be either positive or negative.

**M2b: gBGC hotspots of intensity *B*_1_** = **4*N*_*e*_*b*_1_ in frequency *f* and basal gBGC of intensity ***B*_0_ = **4***N*_*e*_*b***_0_**:

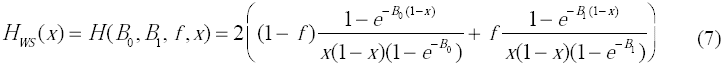

and *H*_SW_(*X*) = *H*(−*B*_0_,−*B*_1_, *f*, *x*). *B*_0_ and *B*_1_ can be either positive or negative.

Polarization errors were included in the four models according to equations (2).

**M3: constrained model of gBGC hotspots**

This model is equivalent to the M2b model, except for that *f* is fixed according to the fraction of recombination hotspots detected by HapMap and *B*_1_ = *ρB*_0_, where *ρ* is the ratio of recombination rates measured in and outside hotspots.

Assuming independence between SNPs, the likelihood of the model can thus be written as:

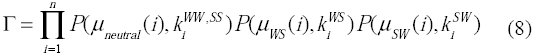

Parameters estimates were obtained by maximization of the log-likelihood function using the FindMaximum function of Mathematica v8 (Wolfram 1996) (R scripts are also available but are slower). The composite parameters *θ*_*WS*_ *θ* = 4*N*_*e*_*uL* and *θ* = 4*N*_*e*_*vL*, *λ*, and the coefficients *r*_*i*_ were constrained to be positive. In model M1, *B* was a free parameter. In model M2a and M2b, *B*_0_ and *B*_1_ were free parameters and *f* was constrained to lie between 0 and 1. Error rates were constrained to be positive and lower than ½. The accuracy and the speed of the maximization are greatly increased by choosing starting values close to the optimum. This is possible because we can obtain rough estimates of most parameters: (i) *θ*, *θ*_*WS*_ are set to the Watterson’s estimates for the corresponding SFS. Watterson’s estimate is also computed for *θ*_*SW*_ to set the mutational bias to *λ* = *θ*_*SW*_ p_*GC*_ /*θ*_*WS*_(1 – *p*_*GC*_). (ii) The initial *r*_*i*_ coefficients are based on the neutral spectrum and set to *r*_*i*_ = *i k_i_* / *k*_1_, where *k*_*i*_ is the number of neutral SNP in frequency *i*/*n*. (iii) To set the initial *B* value in model M1 we used the fact that the log-ratio of the WS and the SW spectra is independent of the *r*_*i*_ coefficients and is linear in *B*:

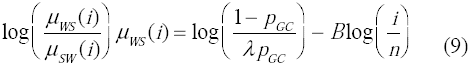

We thus used the slope of the regression of the log-ratio over the log of the class frequencies as the starting value for *B*. (iv) Polarization errors rates are set to 0.01. For models M2a and M2b, parameters obtained by the maximization of model M1 were used as starting values except for the additional parameters *f* set to 0.9. When runs did not converge over long times, other starting values where tested.

Likelihood-ratio tests (LRT) with one degree of freedom can be performed to compare the different nested gBGC models (M1 vs M0, M2a vs M1, M2b vs M2a, with or without polarization errors). Similarly, the equivalent models with and without polarization errors can be compared. Note that because of possible non-independence between SNPs, LRT are anti-conservative and must be viewed with caution. However, maximum likelihood estimates should not be affected by such non-independence.

### Simulations

We simulated datasets by drawing SNPs from Poisson distributions with expectation values given by the population genetics models M0 to M2b. These are the “true” correctly orientated datasets. Then, from these datasets, we built datasets with a given proportion of polarization errors: *e*_neutral_, *e*_WS_, and *e*_SW_. For these “observed” datasets with polarization errors, the observed numbers of SNPs in frequency classes *i*/*n* are thus:

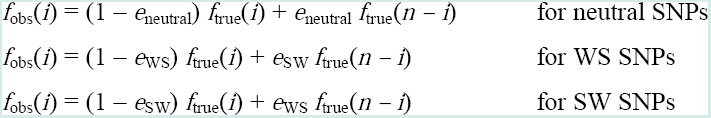

Note that the observed numbers of WS SNPs are proportional to (1 – *p*_*GC*_) θWS and the observed number of SW SNPs to *p*_*GC*_ λ θWS. We then applied the different models, without and with error corrections to the two kinds of datasets. The following parameters are common to all simulations: θneutral = 1000, θWS = 2000, λ = 2, *n* = 20.

## Acknowledgements

This work was supported by the Centre National de la Recherche Scientifique, and by the Agence Nationale de la Recherche (ABS4NGS: ANR-11-BINF-0001-06). This is manuscript ISEM xx.

## References

Abecasis GR, Auton A, Brooks LD, DePristo MA, Durbin RM, Handsaker RE, Kang HM, Marth GT, McVean GA. 2012. An integrated map of genetic variation from 1,092 human genomes. Nature 491(7422): 56–65.

Auton A, Fledel-Alon A, Pfeifer S, Venn O, Segurel L, Street T, Leffler EM, Bowden R, Aneas I, Broxholme J et al. 2012. A fine-scale chimpanzee genetic map from population sequencing. Science 336(6078): 193–198.

Berglund J, Pollard KS, Webster MT. 2009. Hotspots of biased nucleotide substitutions in human genes. PLoS Biol 7(1): e26.

Birdsell JA. 2002. Integrating genomics, bioinformatics, and classical genetics to study the effects of recombination on genome evolution. Mol Biol Evol 19(7): 1181–1197.

Boyko AR, Williamson SH, Indap AR, Degenhardt JD, Hernandez RD, Lohmueller KE, Adams MD, Schmidt S, Sninsky JJ, Sunyaev SR et al. 2008. Assessing the evolutionary impact of amino acid mutations in the human genome. PLoS Genet 4(5):e1000083.

Brown TC, Jiricny J. 1987. A specific mismatch repair event protects mammalian cells from loss of 5-methylcytosine. Cell 50(6): 945–950.

Capra JA, Hubisz MJ, Kostka D, Pollard KS, Siepel A. 2013. A model-based analysis of GC-biased gene conversion in the human and chimpanzee genomes. PLoS Genet 9(8): e1003684.

Charlesworth B. 2013. Why we are not dead one hundred times over. Evolution 67(11): 3354–3361.

Clement Y, Arndt PF. 2013. Meiotic Recombination Strongly Influences GC-Content Evolution in Short Regions in the Mouse Genome. Mol Biol Evol.

Coop G, Myers SR. 2007. Live hot, die young: transmission distortion in recombination hotspots. PLoS Genet 3(3): e35.

De Maio N, Schlotterer C, Kosiol C. 2013. Linking great apes genome evolution across time scales using polymorphism-aware phylogenetic models. Mol Biol Evol 30(10): 2249–2262.

Dreszer TR, Wall GD, Haussler D, Pollard KS. 2007. Biased clustered substitutions in the human genome: the footprints of male-driven biased gene conversion. Genome Res 17(10): 1420–1430.

Duret L, Arndt PF. 2008. The impact of recombination on nucleotide substitutions in the human genome. PLoS Genet 4(5): e1000071.

Duret L, Eyre-Walker A, Galtier N. 2006. A new perspective on isochore evolution. Gene 385: 71–74.

Duret L, Galtier N. 2009. Biased gene conversion and the evolution of mammalian genomic landscapes. Annu Rev Genomics Hum Genet 10: 285–311.

Duret L, Semon M, Piganeau G, Mouchiroud D, Galtier N. 2002. Vanishing GC-rich isochores in mammalian genomes. Genetics 162(4): 1837–1847.

Escobar JS, Glémin S, Galtier N. 2011. GC-Biased Gene Conversion Impacts Ribosomal DNA Evolution in Vertebrates, Angiosperms, and Other Eukaryotes. Mol Biol Evol 28(9): 2561–2575.

Eyre-Walker A. 1998. Problems with parsimony in sequences of biased base composition. J Mol Evol 47(6): 686–690.

Eyre-Walker A, Woolfit M, Phelps T. 2006. The distribution of fitness effects of new deleterious amino acid mutations in humans. Genetics 173(2): 891–900.

Frazer KA Ballinger DG Cox DR Hinds DA Stuve LL Gibbs RA Belmont JW Boudreau A Hardenbol P Leal SM et al. 2007. A second generation human haplotype map of 3.1 million SNPs. Nature 449(7164): 851–861.

Galtier N, Duret L. 2007. Adaptation or biased gene conversion? Extending the nullhypothesis of molecular evolution. Trends Genet 23(6): 273–277.

Galtier N, Duret L, Glémin S, Ranwez V. 2009. GC-biased gene conversion promotes the fixation of deleterious amino acid changes in primates. Trends Genet 25(1): 1–5.

Glémin S. 2010. Surprising fitness consequences of GC-biased gene conversion: I. Mutation load and inbreeding depression. Genetics 185(3): 939–959.

Hernandez RD, Williamson SH, Zhu L, Bustamante CD. 2007. Context-dependent mutation rates may cause spurious signatures of a fixation bias favoring higher GC-content in humans. Mol Biol Evol 24(10): 2196–2202.

Katzman S, Capra JA, Haussler D, Pollard KS. 2011. Ongoing GC-biased evolution is widespread in the human genome and enriched near recombination hotspots. Genome Biol Evol.

Keightley PD, Eyre-Walker A. 2007. Joint inference of the distribution of fitness effects of deleterious mutations and population demography based on nucleotide polymorphism frequencies. Genetics 177(4): 2251–2261.

Kong A, Frigge ML, Masson G, Besenbacher S, Sulem P, Magnusson G, Gudjonsson SA, Sigurdsson A, Jonasdottir A, Jonasdottir A et al. 2012. Rate of de novo mutations and the importance of father’s age to disease risk. Nature 488(7412): 471–475.

Kostka D, Hubisz MJ, Siepel A, Pollard KS. 2012. The role of GC-biased gene conversion in shaping the fastest evolving regions of the human genome. Mol Biol Evol 29(3): 1047–1057.

Lachance J, Tishkoff SA. 2014. Biased gene conversion skews allele frequencies in human populations, increasing the disease burden of recessive alleles. Am J Hum Genet 95(4):408–420.

Langergraber KE, Prufer K, Rowney C, Boesch C, Crockford C, Fawcett K, Inoue E, Inoue-Muruyama M, Mitani JC, Muller MN et al. 2012. Generation times in wild chimpanzees and gorillas suggest earlier divergence times in great ape and human evolution. PNAS 109(39): 15716–15721.

Lartillot N. 2013a. Interaction between selection and biased gene conversion in mammalian protein-coding sequence evolution revealed by a phylogenetic covariance analysis. Mol Biol Evol 30(2): 356–368.

Lartillot N.. 2013b. Phylogenetic patterns of GC-biased gene conversion in placental mammals and the evolutionary dynamics of recombination landscapes. Mol Biol Evol 30(3): 489–502.

Lesecque Y, Glémin S, Lartillot N, Duret L. 2014. The Red Queen Model of Recombination Hotspots Evolution in the Light of Archaic and Modern Human Genomes. PLoS Genetics **In press**.

Lesecque Y, Keightley PD, Eyre-Walker A. 2012. A resolution of the mutation load paradox in humans. Genetics 191(4): 1321–1330.

Lesecque Y, Mouchiroud D, Duret L. 2013. GC-biased gene conversion in yeast is specifically associated with crossovers: molecular mechanisms and evolutionary significance. Mol Biol Evol 30(6): 1409–1419.

Marais G. 2003. Biased gene conversion: implications for genome and sex evolution. Trends Genet 19(6): 330–338.

Muyle A, Serres-Giardi L, Ressayre A, Escobar J, Glémin S. 2011. GC-biased gene conversion and selection affect GC content in the *Oryza* genus (rice). Mol Biol Evol 28(9): 2695–2706.

Myers S, Bottolo L, Freeman C, McVean G, Donnelly P. 2005. A fine-scale map of recombination rates and hotspots across the human genome. Science 310(5746): 321–324.

Nagylaki T. 1983. Evolution of a finite population under gene conversion. Proc Natl Acad Sci USA 80(20): 6278–6281.

Necsulea A, Popa A, Cooper DN, Stenson PD, Mouchiroud D, Gautier C, Duret L. 2011. Meiotic recombination favors the spreading of deleterious mutations in human populations. Hum Mutat 32(2): 198–206.

Paten B, Herrero J, Fitzgerald S, Beal K, Flicek P, Holmes I, Birney E. 2008. Genome-wide nucleotide-level mammalian ancestor reconstruction. Genome Res 18(11): 1829–1843.

Pessia E, Popa A, Mousset S, Rezvoy C, Duret L, Marais GA. 2012. Evidence for widespread GC-biased gene conversion in eukaryotes. Genome Biol Evol 4(7): 675–682.

Ptak SE, Hinds DA, Koehler K, Nickel B, Patil N, Ballinger DG, Przeworski M, Frazer KA, Paabo S. 2005. Fine-scale recombination patterns differ between chimpanzees and humans. Nat Genet 37(4): 429–434.

Ratnakumar A, Mousset S, Glémin S, Berglund J, Galtier N, Duret L, Webster MT. 2010. Detecting positive selection within genomes: the problem of biased gene conversion. Phil Trans R Soc Lond B 365(1552): 2571–2580.

Romiguier J, Ranwez V, Douzery EJ, Galtier N. 2010. Contrasting GC-content dynamics across 33 mammalian genomes: relationship with life-history traits and chromosome sizes. Genome Res 20(8): 1001–1009.

Serres-Giardi L, Belkhir K, David J, Glémin S. 2012. Patterns and evolution of nucleotide landscapes in seed plants. Plant Cell 24(4): 1379–1397.

Sjolund AB, Senejani AG, Sweasy JB. 2013. MBD4 and TDG: multifaceted DNA glycosylases with ever expanding biological roles. Mutat Res 743-744: 12–25.

Smith NG, Eyre-Walker A. 2001. Synonymous codon bias is not caused by mutation bias in G+C-rich genes in humans. Mol Biol Evol 18(6): 982–986.

Spencer CC, Deloukas P, Hunt S, Mullikin J, Myers S, Silverman B, Donnelly P, Bentley D, McVean G. 2006. The influence of recombination on human genetic diversity. PLoS Genet 2(9): e148.

Surtees JA, Argueso JL, Alani E. 2004. Mismatch repair proteins: key regulators of genetic recombination. Cytogenet Genome Res 107(3-4): 146–159.

Webster MT, Axelsson E, Ellegren H. 2006. Strong regional biases in nucleotide substitution in the chicken genome. Mol Biol Evol 23(6): 1203–1216.

Winckler W, Myers SR, Richter DJ, Onofrio RC, McDonald GJ, Bontrop RE, McVean GA, Gabriel SB, Reich D, Donnelly P et al. 2005. Comparison of fine-scale recombination rates in humans and chimpanzees. Science 308(5718): 107–111.

Wolfram S. 1996. The Mathematica book. Cambridge University Press, Cambridge.

